# Can paid reviews promote scientific quality and offer novel career perspectives for young scientists?

**DOI:** 10.1101/089490

**Authors:** Christian Wurzbacher, Hans-Peter Grossart, Erik Kristiansson, R. Henrik Nilsson, Martin Unterseher

## Abstract

We exemplify how community driven paid reviews could work in conjunction with a feed-back loop to young scientists. This would remove pressure from the conventional peer-review system and promote the integration of reviews into an academic career. This concept may eventually revive the community based journal type.

## Introduction

The standard system to secure quality and error correction of scientific publications is peer review. The peer review system is, however, currently under high pressure due to the rapid increase in the number of scientific publications globally, with a growth rate of 9% per year [1]. The scientific publishing field is characterised by many recent changes, such as the emergence of fast publication open access (OA) journals and the increasing use of preprint servers such as bioRxiv.org to counter long processing times. This ever-increasing pressure is also manifested in the publishing process itself and facilitates a decreasing trust in scientific integrity, a phenomenon called “overflow” [3]. Examples for these concerns can be find on all different levels of the publishing process: a) on the journal level, where so called “predatory journals” impose article processing charges (APC) without a credible peer-review process and where many journals have difficulties to find qualified editors to handle the flood of submitted papers; b) on the editor level, where editors struggle to find adequate reviewers [3]; and often turn to the manuscript authors for reviewer suggestions; c) on the author level, where many authors complain about long waiting times, superficial reviews, and non-responding editors; and d) on the reviewer level, where many scientists receive an increasing number of review requests during the progression of their career, adding to their significant time-constraints and duties as, e.g., supervisors, scientists, and lecturers. In this opinion piece we focus on the reviewer level, since reviewers represent the backbone of the whole scientific peer-review system. It is also the level where we feel that change is the easiest to bring about.

### The reviewer dilemma

The backbone of the peer-review system is based on honour, reliability, and trustworthiness. An implicit assumption is that every scientist will accept offers to review manuscripts and applications as long as these fall within the expertise of the scientist. This unspoken duty is based on a give-and-take approach within the community – a scientist is implicitly expected to review at least as many manuscripts as s/he submits for publication annually - such that a sort of global workload balance is attained. Although serving as a reviewer is indeed seen as a community service by the scientific community [2], this service is usually not stipulated in the working contract of scientists. Therefore, exactly how much time the researcher should invest in this community service is each scientist’s personal decision. The fact that this is up to each individual researcher to decide may already raise questions, but what really makes this community service disputable is the fact that publishers make profit from it. Publishing houses charge either publication fees from authors or institutions, or article access fees from readers for accessing published research. At the same time, the underlying articles were processed with free-of-charge review services – services that were already paid for once in form of the researcher’s salary (often tax money). A second critical argument is that this community service is not visible to other scientists, because it is typically an anonymous process. Therefore, the contributions of even very active individuals are not valued per se, since their work is to a large extend unnoticed (sometimes even undesired, in case of rejections). Some publishers have already recognised these problems and suggested various changes to the system, such as open, credited review processes or the rewarding of reviewers with increased attention, gifts, or financial waives on APCs. In addition, community-driven initiatives such as Publons (http://prw.publons.com/) have emerged to increase the visibility and reputation of reviewers. Still, these efforts can be only considered as a temporary solution and as gestures that will not solve these critiques in a sustainable manner.

The authors of this manuscript are researchers, article authors, editors, and frequent reviewers, and we have experienced the shortcomings of the present system from the respective points of view. We argue that the situation is untenable in the long run, and in the present opinion piece we discuss an alternative where reviewers are honoured by payment. This, we argue, will raise reviews from an anonymous community service to a visible and quantifiable service. Even if the review process stays anonymous, this service will offer a new job profile, in particular for postdocs, that can be integrated in a regular scientific career and serve to strengthen the weight of future job or research grant applications of these individuals.

## Gedankenexperiment: careful implementation of paid reviews into a journal’s concept

The introduction of paid reviews into the current peer-review system would have far-reaching implications and impacts on all levels of scientific publishing, ranging from reviewers (most obviously) through authors and editors to journals. Here we exemplify paid reviews through a novel fictive journal concept built on this idea. Paid reviews could be also operated in other ways, but for the purpose of the present opinion piece, a fictive journal will help to illustrate these major changes.

### Potential financial implications

During the last 10 years, the publishing sector has undergone major changes, notably through the introduction of OA journals. Beside community-driven OA journals (e.g. the Pensoft journals, eLife until 2016), the majority of the commercial publishers use a business model that implements APC. These APCs usually range from 200-3000 e, depending on the journal and the level of service provided (see [5]). For instance, the APC for a PLOS ONE article is currently 1330 € (October 2016), however, the costs for the publisher to process and publish the article are much less, approximately 260 € [5]. Our fictive journal is OA since it (1) readily allows us to highlight the immediate effects of paid reviews on the financial balance of the journal, and (2) it promotes the most immediate and effective scientific transfer. The former is of course the major issue when we start paying for reviews. We assume the following: 1) a thorough review made by an expert takes on average 4-8 hours of work depending on several factors such as the experience of the reviewer in the field, the length of the manuscript, the number of references, and the complexity of the presented ideas; 2) a reviewer will cost between 40-150 € per hour, mainly depending on the country of residence; and 3) we would like to make use of three independent reviewers per submitted manuscript (two reviews are often considered as sufficient). None of these assumptions go against common practice in their respective field of application.

Suppose a manuscript was submitted to our fictive journal. The manuscript consumed an average of, say, 5 hours of work for each of the three reviewers, and the reviewers were paid on average 100 € per hour. The review cost will thus be around 1500 €. Together with the handling and publishing costs, we will tangent the present upper range (3000 €) for current OA journal APCs without considering profit. On top of that, accepted articles will have to compensate for the costs for any rejected – but still reviewed – manuscripts. This additional cost presents a problem to us, and we decided that the one realistic way to deal with submitted-but-rejected manuscripts is to charge the authors an initial APC of 500 € at the time of submission, without guaranteeing any article acceptance. We justify this by arguing that the journal provides a service (it administers and pays the reviewers, and the resulting reviews will always be sent to the authors upon completion of the review process) and this has to be paid for, much in the same way as you pay for professional article editing services. Although this hurdle looks tough, it will have two immediate positive effects: (1) it will prevent the majority of premature or fragmentary manuscripts from being submitted in the first place “just in case it gets accepted”, and (2) authors who do choose to submit their manuscript to the journal are likely to double-check all aspects of their manuscript to make sure that the 500 € are invested well. In summary, we think that the initial 500 € APC will lead to fewer, but better, submissions. It furthermore puts significant pressure on the reviewers to do a thorough, timely job and to provide detailed, in-depth feedback – which is not always the case in the present system. With this practice we hope to increase the manuscript acceptance rate to above 90% - for comparison, PLOS ONE has an acceptance rate of 70% [5]. We speculate that a high acceptance rate will eventually fully finance the reviews of rejected manuscripts and will compensate for reduced APCs for low-income countries. These pre-charges can be also seen critical and as an alternative concept we can imagine to install a hybrid review format with a mixture of conventional and paid reviewers that is supported by external funding.

Another advantage, especially for young scientists who are in need to publish timely, is that such a journal concept could actually promise short and well-defined turn-around times if needed (e.g., of 14 days) based on a fixed agreement with the reviewers.

### The reviewers and former editors

In order to ensure reviews of high quality, we will aim to hire experienced researchers who have published a certain number of articles (e.g., more than 15 articles, including at least 3 first/corresponding authorships, during the last 5 years) in various international, ISI-covered peer-reviewed journals. Our main target reviewer group will be postdoctoral researchers – ideally those having completed at least two postdoc terms – and who struggle to find a faculty position. This is the time point in academia where most young scientists who still have a passion for science are forced to drop out of science due to lack of future prospects (those lacking a passion for science would, by and large, have left already after the first postdoctoral term). This “lost generation” is one of the most pressing problems of a scientific career today and was titled as “the academic research crisis” [4]. Thus, we explicitly aim for very experienced but unemployed or part-time working (young) scientists, and we advice not to pay for a review from a full-time employed scientist. It is our hope that paid reviews will offer a temporary career alternative in the academic working environment. For instance, postdocs can work half-time as reviewers and half-time on the next grant proposal. Alternatively, the postdocs could spend a couple of years doing full-time paid reviews and integrate this into their CVs. Several flexible working solutions can emerge that finally might allow for reintegration of young scientists back into science. This is an important point, because unlike in conventional publishing with a private publisher, the APC money will be reinvested into the scientific community. On top of that it might help to reduce the current discrepancy between number of active authors and number of active reviewers [3].

Another issue that came up during our discussions is that editors may lose some of their importance in this concept. The implementation of paid reviews after paying an initial APC renders an initial screening of the manuscript by an editor somewhat obsolete: the manuscript will always enter the review process. Similarly, the discussions on the pros and cons of a manuscript and the final decision can be made by the three highly educated reviewers, who basically will replace the role of an editorial board. We recognize that this is at odds with how most journals are run now, but we do not necessarily see this as a bad thing.

### Additional considerations

Our concept is probably not applicable to all types of articles. For example, field or laboratory method development papers may be difficult to review for reviewers who do not, or no longer, have direct access to specific scientific equipment. Similarly, scientific topics that are overly specific or highly theoretical may be difficult to address with a limited pool of reviewers, experienced as they may be. The journal will have to outline specifically what type of manuscripts it will be able to review. This outline will always depend on the pool of available reviewers and is thus subjected to change over the course of time. Moreover, we would like to emphasise that we do not propose to change the peer-review system *per se*, rather offer alternative conditions under which it can operate. We are convinced that this will help to increase the quality of peer-review above the current high standards.

### Community journal

It does seem problematic to combine our approach with substantial profits, as the APCs would run very high. We thus feel that our approach is best suited for community-driven journals or by publishing companies with a strong passion for the underlying science, rather than for commercial, large-scale publishing houses. Thus, whether journals adopting our approach will ever be able to compete – in terms of impact factor or otherwise – with leading journals from leading publishing houses is unclear. Next to substantial financial support, a journal using paid reviewers requires some degree of acceptance in the scientific community, which is willing to promote the career of young scientists in the field.

## Summary

We present a paid peer-review concept that promises fixed article processing times. Unlike the situation with other contemporary journals, the article charges are redirected to the scientific community, especially to the “lost generation” of highly qualified postdoctoral researches who have been unable to secure a faculty position so far. This community-oriented peer-review concept involves additional costs but in turn promises a high standard of scientific endeavors and a feedback loop to the scientific community. We do think that our model offers something that contemporary journals struggle to provide: highly motivated reviewers, who lack a hidden agenda at that. We do not anticipate that our model will go down well with large-scale commercial models for scientific publishing, and we question whether large-scale commercial models for scientific publishing indeed are in the best interest of the scientific community in the first place.

## Open Feedback

Finally, we hereby open this article for your feedback. We have installed a poll: https://goo.gl/ZbTKnM and will publish the results with the next revision of the article in January 2017. We are encouraging scientists from all fields covered by bioRxiv.org to comment and contribute. We are furthermore willing to acknowledge you in the manuscript or to add you to the author list if you can make a significant conceptual contribution to this discussion and to the manuscript.

